# Disease-modifying effects of natural Δ9-tetrahydrocannabinol in endometriosis-associated pain

**DOI:** 10.1101/715938

**Authors:** Alejandra Escudero-Lara, Josep Argerich, David Cabañero, Rafael Maldonado

**Affiliations:** Laboratory of Neuropharmacology, Department of Experimental and Health Sciences, Universitat Pompeu Fabra. Barcelona, Spain; IMIM (Hospital del Mar Medical Research Institute), Barcelona, Spain

## Abstract

Endometriosis is a chronic painful disease highly prevalent in women that is defined by growth of endometrial tissue outside the uterine cavity and lacks adequate treatment. Medical use of cannabis derivatives is a current hot topic and it is unknown whether phytocannabinoids may modify endometriosis symptoms and development. Here we evaluate the effects of repeated exposure to Δ9-tetrahydrocannabinol (THC) in a mouse model of surgically-induced endometriosis. In this model, female mice develop pelvic mechanical hypersensitivity, anxiety-like behavior and sharp memory deficits associated to the presence of extrauterine endometrial cysts. Interestingly, chronic THC (2 mg/kg/day) provides sustained alleviation of pelvic hypersensitivity without altering the anxiogenic phenotype, modifies uterine innervation and restores cognitive function, an effect correlated with neuroinflammatory changes in prefrontal cortex. Strikingly, THC also inhibits the development of endometrial cysts. These data highlight the interest of scheduled clinical trials designed to investigate possible benefits of THC for women with endometriosis.

## Introduction

Endometriosis is a chronic inflammatory disease that affects 1 in 10 women in childbearing age (Zondervan KT, Becker CM, Koga K, Missmer SA, Taylor RN, 2019). It is characterized by the growth of endometrium in extrauterine locations, chronic pelvic pain, infertility, emotional distress and loss of working ability (Fourquet, Báez, Figueroa, Iriarte, & Flores, 2011; Márki, Bokor, Rigó, & Rigó, 2017; Zondervan KT, Becker CM, Koga K, Missmer SA, Taylor RN, 2019). Current clinical management provides unsatisfactory outcomes. On the one hand, hormonal therapy has unwanted effects including contraception and emotional disturbances (Ross & Kaiser, 2017; Skovlund, Mørch, Kessing, & Lidegaard, 2016), whereas surgical excision of the growths is associated with high-recurrence rates and post-surgical pain (Garry, 2004). Hence, clinical treatments are limited and women often unsatisfactorily self-manage their pain (Armour, Sinclair, Chalmers, & Smith, 2019). In this context, marijuana legalization for medical purposes in American and European states has led to increased availability of phytocannabinoids (Abuhasira, Shbiro, & Landschaft, 2018). While cannabis may provide pain relief in certain conditions (Campbell et al., 2001), it is unclear whether it may modify endometriosis symptoms or development.

Δ9-tetrahydrocannabinol (THC) is the main psychoactive constituent of the *Cannabis sativa* plant, and multiple animal and clinical studies suggest that could be effective relieving chronic pain (De Vry, Kuhl, Franken-Kunkel, & Eckel, 2004; Harris, Sufka, Gul, & Elsohly, 2016; King et al., 2017; Ueberall, Essner, & Mueller-Schwefe, 2019; Williams, Haller, Stevens, & Welch, 2008), although controversial results have been obtained in human clinical trials (Stockings et al., 2018). However, THC has important side effects including cognitive deficits and anxiety (Célérier et al., 2006; Kasten, Zhang, & Boehm II, 2017; Puighermanal et al., 2013). This work investigates the effects of natural THC in a mouse model of endometriosis that mimics the clinical condition of women. Our data show that THC is effective inhibiting pelvic hypersensitivity and the abrupt cognitive impairment observed in mice with ectopic endometrium. Interestingly, these effects correlate with neuroplastic changes in peripheral innervation and inflammatory modifications in prefrontal cortex. Notably, THC also shows efficacy limiting the development of the ectopic endometrium, revealing disease-modifying effects of this natural cannabinoid.

## Results and discussion

### Ectopic endometrium leads to mechanical hypersensitivity, anxiety-like behavior and memory impairment

Our first aim was to characterize a novel experimental procedure to evaluate at the same time nociceptive, cognitive and emotional manifestations of endometriosis in female mice. Mice were subjected to a surgical implantation of endometrial tissue in the peritoneal wall of the pelvic compartment or to a sham procedure. Mice receiving ectopic endometrial implants developed persistent mechanical hypersensitivity in the pelvic area, whereas sham mice recovered their baseline sensitivity and showed significant differences in comparison to endometriosis mice since the second week of implantation (Figure 1a and Figure 1 – figure supplement 1). Endometriosis mice also showed anxiety-like behavior revealed by increased percentage of time and entries to the closed arms of an elevated plus maze (Figure 1b). In line with these findings, previous rodent models of endometriosis found increased mechanosensitivity in the lower abdomen (Arosh et al., 2015; Greaves et al., 2017) and affective-like disturbances (Filho et al., 2019; Li et al., 2018), although this cognitive impairment has not yet been revealed in rodent models of endometriosis. Previous works associate nociceptive and emotional distress in chronic pain settings with cognitive decline (Bushnell et al., 2015; La Porta et al., 2015; You et al., 2018). We revealed in our model a dramatic impairment of long-term memory in endometriosis mice (Figure 1c). While mnemonic effects of this pathology have not been thoroughly evaluated, a cognitive impairment may contribute to the loss of working ability consistently reported in women with endometriosis (Hansen, Kesmodel, Baldursson, Schultz, & Forman, 2013; Sperschneider et al., 2019). Hence, mice with ectopic endometrium recapitulate in our model the main symptomatology observed in the clinics. Mice receiving endometrial implants developed 3 to 5 endometrial cysts in the peritoneal wall of the pelvic compartment. Cysts were of 2.59 ± 0.34 mm diameter, filled of fluid, with glandular epithelium and stroma and innervated by beta-III tubulin positive fibers (Figure 1d), as shown in women (Tokushige, Markham, Russell, & Fraser, 2006; Wang, Tokushige, Markham, & Fraser, 2009) and other rodent models (Arosh et al., 2015; Berkley, Dmitrieva, Curtis, & Papka, 2004). Interestingly, we also found uterine hyperinnervation in endometriosis mice (Figure 1 – figure supplement 2), reproducing not only the symptoms, but also the histological phenotype observed in women with endometriosis (Miller EJ, 2015; Tokushige et al., 2006).

**Figure 1.**
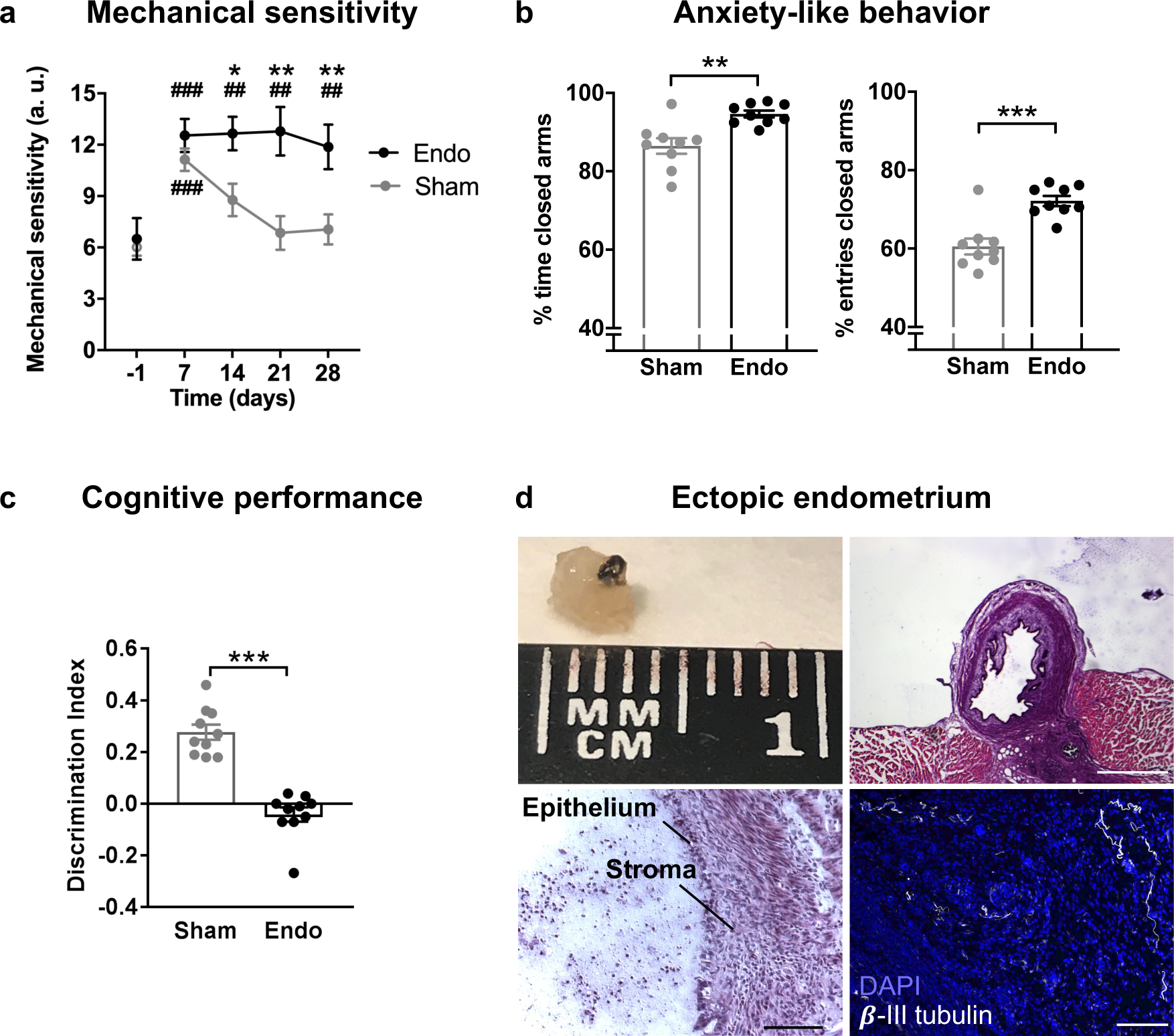
Behavioral and histological alterations in female mice with ectopic endometrial implants. Endometriosis mice showed (**a**) persistent increase in pelvic mechanical sensitivity (**b**) increased anxiety-like behavior and (**c**) cognitive impairment. (**d**) Cysts recovered from endometriosis mice were filled of fluid, contained endometrial epithelium and stroma and were innervated by beta-III tubulin-labeled fibers. Blue is DAPI and white is β-III tubulin. Scale bar = 1 mm (upper right), 100 μm (lower left and lower right). Error bars are mean ± SEM. One-way repeated measures ANOVA + Bonferroni (a) and Student t-test (b and c). *p<0.05, **p<0.01, ***p<0.001 vs sham. ##p<0.01, ###p<0.001 vs baseline. Endo, endometriosis; a.u., arbitrary units. Figure 1 - figure supplements 1 and 2. Figure 1 - Source Data.

### Δ9-tetrahydrocannabinol alleviates pelvic pain, restores cognitive function and limits the growth of ectopic endometrium

Our second objective was to assess the effects of THC exposure on the endometriosis model to select an appropriate dose for a chronic treatment. Acute doses of THC were first tested in endometriosis and sham mice. Acute THC administration produced a dose-dependent reduction of pelvic mechanical hypersensitivity (Figure 2). The acute ED50 of THC 1.916 mg/kg (≈2 mg/kg) was chosen for the repeated administration.

**Figure 2.**
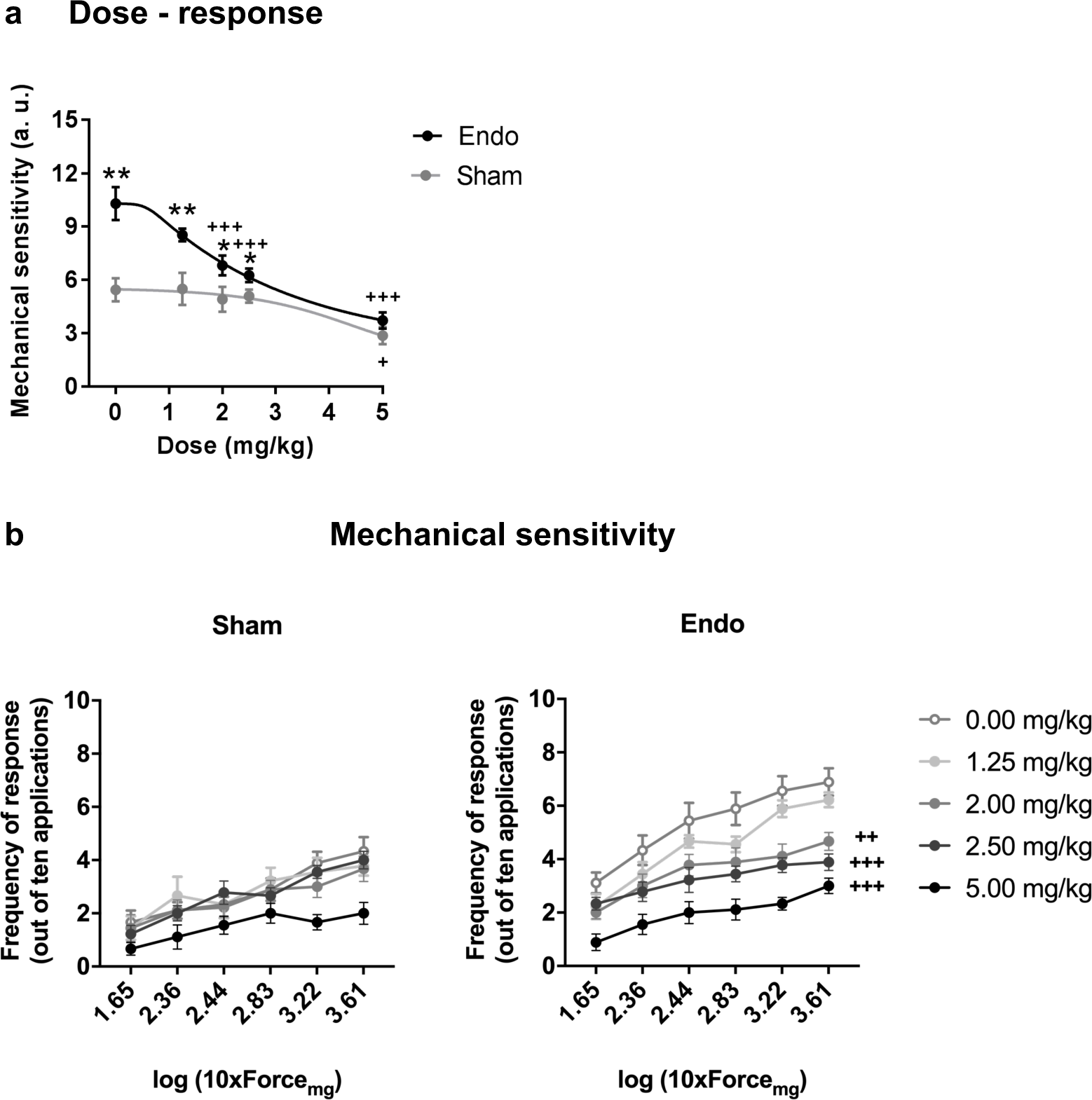
Effect of acute THC administration on the nociceptive responses to mechanical stimulation. (**a**) Acute THC produced a dose-dependent reduction of pelvic mechanical hypersensitivity. (**b**) Administration of 2, 2.5 and 5 mg/kg of THC alleviated mechanical sensitivity in endometriosis mice. Error bars are mean ± SEM. One-way repeated measures ANOVA + Bonferroni. *p<0.05, **p<0.01 vs sham; +p<0.05, ++p<0.01, +++p<0.001 vs vehicle. Endo, endometriosis; THC, Δ9-tetrahydrocannabinol; a.u., arbitrary units. Figure 2 - Source Data.

Repeated exposure to THC 2 mg/kg, once daily for 32 days, provided a sustained alleviation of mechanical pain during the whole treatment period (Figure 3a and Figure 3 - figure supplement 1) without inducing tolerance. The absence of tolerance to THC-induced antinociception is in contrast with the tolerance described at higher THC doses in other pain models (Greene, Wiley, Yu, Clowers, & Craft, 2018; Lafleur, Wilson, Morgan, & Henderson-Redmond, 2018; Wakley, Wiley, & Craft, 2014). Our experimental conditions showed no significant effects of repeated THC on anxiety-like behavior in sham or endometriosis mice (Figure 3b). Acute exposure to high THC doses (>7.5 mg/kg) is linked to anxiety-like responses in male mice (Célérier et al., 2006; Puighermanal et al., 2013), whereas low doses (<1,5 mg/kg) produce anxiolytic effects (Célérier et al., 2006; Rubino et al., 2008). Hence, the intermediate dose used in our study did not produce significant modifications of anxiety-like behavior in female mice. Since nociceptive manifestations were abolished in endometriosis mice, these data suggest that alleviation of mechanical pain is independent of anxiety-like responses.

**Figure 3.**
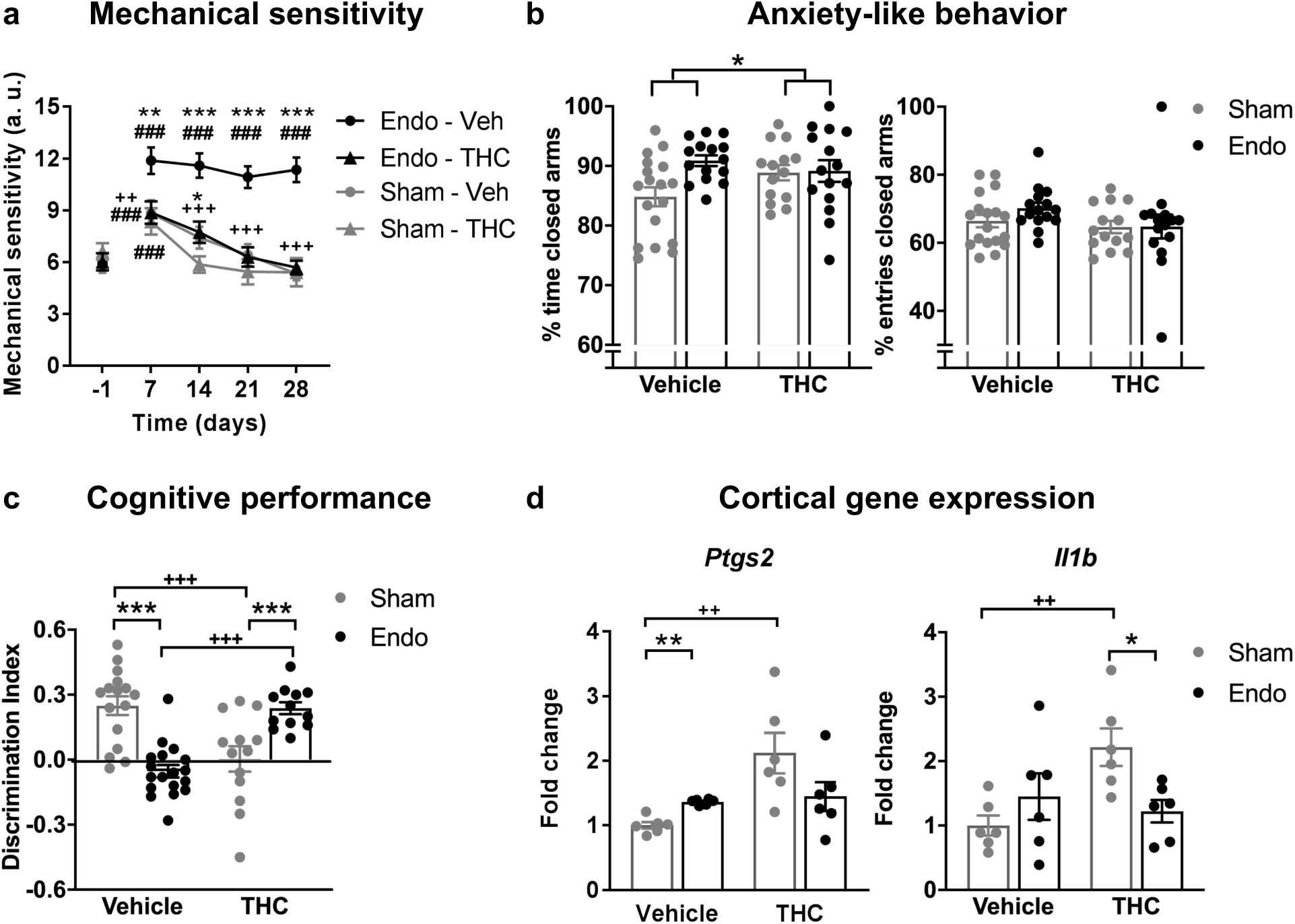
Effects of THC on the behavioral and gene expression changes observed in mice with ectopic endometrium. Repeated THC (**a**) alleviated mechanical hypersensitivity of endometriosis mice but (**b**) did not modify anxiety-like behavior. (**c**) While sham mice receiving THC showed memory impairment, endometriosis mice improved cognitive performance after THC. (**d**) Memory impairment correlated with expression levels of the genes coding for cyclooxygenase-2 (*Ptg2*) and interleukin 1-beta (*Il1b*). Error bars are mean ± SEM. Two-way repeated measures ANOVA + Bonferroni (a), two-way ANOVA + Bonferroni (b and c), and Kruskal-Wallis + Mann Whitney U (d). ###p<0.001 vs baseline. *p<0.05, **p<0.01, ***p<0.001 vs sham. ++p<0.01, +++p<0.001 vs vehicle. Endo, endometriosis; THC, Δ9-tetrahydrocannabinol; a.u., arbitrary units. Figure 3 - figure supplements 1 and 2. Figure 3 - Source Data.

Memory performance was also assessed the third week after starting the THC treatment. As expected, mice exposed to the chronic nociceptive manifestations of endometriosis showed a pronounced cognitive impairment, as well as sham mice exposed to THC, in accordance with previous reports in naïve males (Kasten et al., 2017; Puighermanal et al., 2013). Surprisingly, endometriosis mice repeatedly treated with natural THC showed intact discrimination indices (Figure 3c) suggesting protective effects of THC in this chronic inflammatory condition. In agreement, recent studies have shown cognitive improvements after THC exposure in old male and female mice (Bilkei-Gorzo et al., 2017; Sasson, Rachmany, Toledano, Sarne, & Doron, 2017). These studies suggest an effect of THC in brain areas related to cognitive function, such as prefrontal cortex and hippocampus (Bilkei-Gorzo et al., 2017; Sasson et al., 2017). Thus, we explored whether the cognitive impairment observed in our experimental conditions (Figure 3c) could be accompanied by neuroinflammatory alterations in these brain areas (Figure 3d). We found that cognitive deficits of endometriosis mice treated with vehicle and sham mice receiving THC were associated with increased expression of the genes coding for cyclooxygenase-2 (*Ptg2*) and interleukin 1-beta (*Il1b*) in the medial prefrontal cortex (Figure 3d). However, these changes were not observed in the hippocampus (Figure 3 - supplement 2). Remarkably, the effects of THC were significantly different in endometriosis and sham mice (Figure 3c, 3d). Both aging and chronic pain are associated to local inflammatory events in these brain areas (Di Benedetto, Müller, Wenger, Düzel, & Pawelec, 2017; Ong, Stohler, & Herr, 2019). Our data suggest that these neuroinflammatory conditions modify the effects of THC on cognitive function, allowing to reveal an improvement of memory and inflammatory markers after chronic THC treatment.

Exogenous and endogenous cannabinoids have shown modulatory effects on the female reproductive system (Walker, Holloway, & Raha, 2019). Thus, we analyzed the effects of THC on the ectopic and eutopic endometrium and on ovarian follicle maturation. Interestingly, endometriosis mice receiving chronic THC showed an evident reduction in the size of endometrial cysts (cyst diameter and area of endometrial tissue) without significant effects on cyst innervation (Figure 4 – figure supplement 1a). In agreement, a previous study showed antiproliferative effects of WIN 55212-2, a synthetic cannabinoid agonist, on endometrial cell cultures and in ectopic endometrium implanted to immunodepressed mice (Leconte et al., 2010). The assessment of the uterine diameter and the area of eutopic endometrium (Figure 4 – figure supplement 1b) showed no effects of the THC treatment, suggesting that the antiproliferative activity of THC is restricted to the ectopic endometrium. On the contrary, THC produced an increase of uterine innervation in sham mice, similar to the increase provoked by the ectopic endometrium (Figure 4b) Interestingly, THC prevented uterine hyperinnervation in endometriosis mice suggesting again that THC exposure may have different consequences under chronic inflammatory conditions. We also assessed possible effects of THC on ovarian functioning, since previous works have suggested inhibitory effects of THC on folliculogenesis and ovulation (Adashi, Jones, & Hsueh, 1983; Konje et al., 2009). However, numbers of preantral follicles, antral follicles and corpora lutea remained unchanged in our experimental conditions regardless of the presence of ectopic endometrium or THC (Figure 4 – figure supplement 1c).

**Figure 4.**
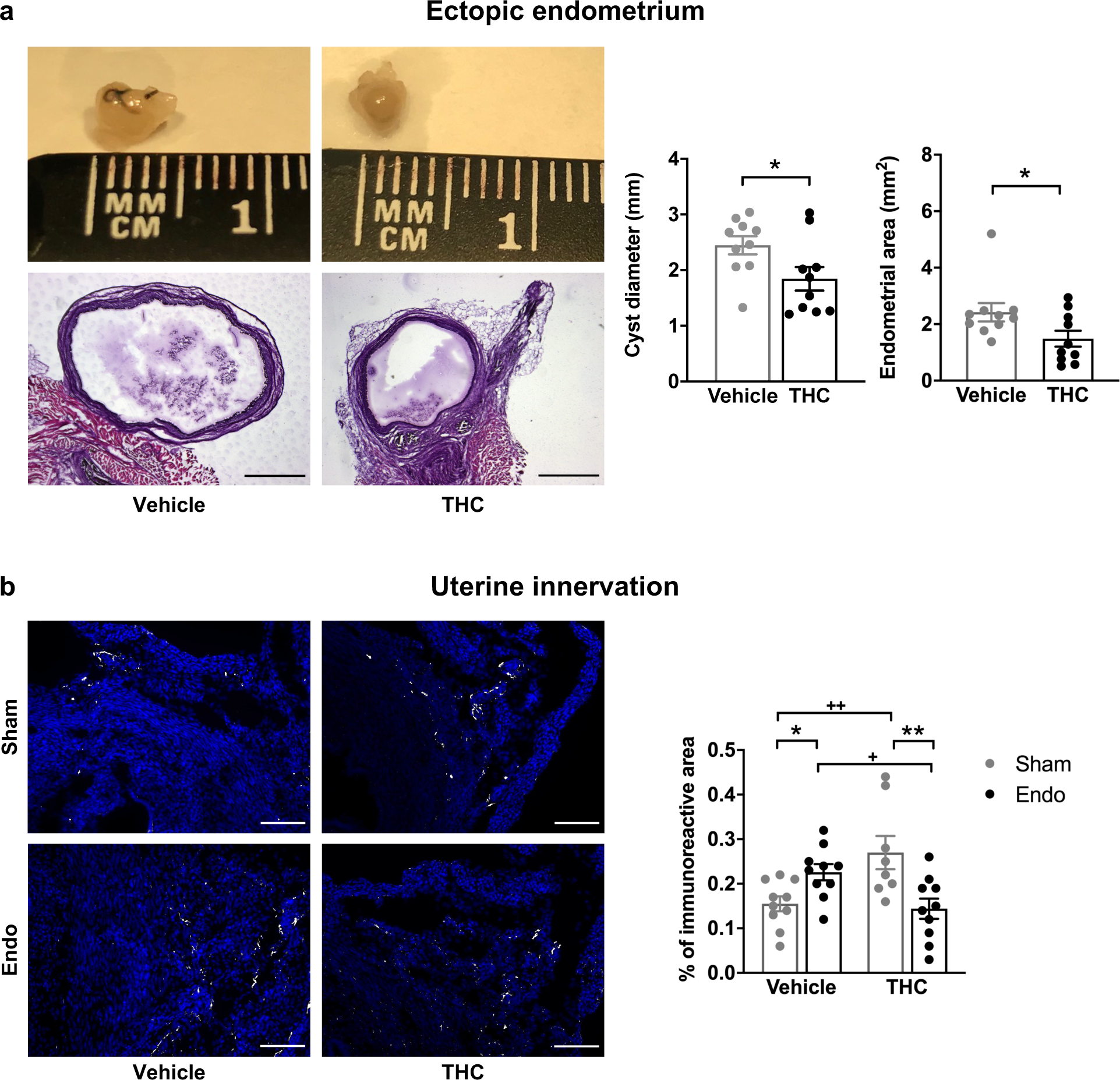
Effects of THC on the histological changes observed in mice with ectopic endometrium. (**a**) Ectopic endometrial growths of mice treated with THC were smaller and had less endometrial tissue than those of mice receiving vehicle. Scale bar = 1 mm. (**b**) THC increased innervation in sham mice but prevented uterine hyperinnervation in endometriosis mice. Blue is DAPI and white is β-III tubulin. Scale bar = 100 μm. Error bars are mean ± SEM. Student t-test (a) and two-way ANOVA + Bonferroni (b). *p<0.05, **p<0.01 vs sham. +p<0.05, ++p<0.01 vs vehicle. Endo, endometriosis; THC, Δ9-tetrahydrocannabinol; a.u., arbitrary units. Figure 4 - Figure supplement 1. Figure 4 - Source data.

## Conclusions

Here we show for the first time that chronic administration of a moderate dose of the phytocannabinoid THC relieves pain sensitivity and cognitive impairment associated to the presence of ectopic endometrial cysts. These behavioral manifestations correlate with a decrease in the size of ectopic endometrium in THC-exposed mice. Pain relief without modification of anxiety-like behavior suggests that anxiety and pain are independent events elicited by the presence of ectopic endometrium. Interestingly, THC produced opposite cognitive effects in sham and endometriosis mice. THC induced cortical expression of neuroinflammatory markers and uterine hyperinnervation in sham animals, but reduced such neuroinflammatory changes in endometriosis mice and prevented uterine hyperinnervation, suggesting different effects of THC under chronic inflammatory conditions. Importantly, THC also inhibited the growth of ectopic endometrium without apparent consequences on the female reproductive system. Altogether, the present data underline the interest of conducting clinical research to assess the effects of moderate doses of THC on endometriosis in patients. Based on our results, we (clinicaltrials.gov, #NCT03875261) and others (gynica.com) have planned the initiation of clinical trials to provide evidence on the translatability of these results to women with endometriosis. These novel clinical trials will be achieved in the following years, allowing to evaluate this new possible endometriosis treatment under pathological human conditions.

## Materials and methods

### Animals

Female C57Bl/6J mice (Charles Rivers, Lyon, France) were used in all the experiments. Mice were 8 weeks old at the beginning of the experiments and were housed in cages of 4 to 5 mice with *ad libitum* access to water and food. The housing conditions were maintained at 21 ± 1°C and 55 ± 10% relative humidity in controlled light/dark cycle (light on between 8 AM and 8 PM). Animals were habituated to housing conditions and handled for 1 week before the start of the experiments. All animal procedures were conducted in accordance with standard ethical guidelines (European Communities Directive 2010/63/EU and NIH Guide for Care and Use of Laboratory Animals, 8^th^ Edition) and approved by autonomic (Generalitat de Catalunya, Departament de Territori i Sostenibilitat) and local (Comitè Ètic d’Experimentació Animal, CEEA-PRBB) ethical committees. All experiments were performed blinded for pharmacological and surgical conditions, with treatments randomized between groups.

### Drugs

Natural THC (THC-Pharm-GmbH, Frankfurt, Germany) was diluted in a vehicle composed of 2.5% ethanol, 5% Cremophor EL (C5135, Sigma-Aldrich St. Louis, MO, USA), and 92.5% saline, and was administered subcutaneously in a volume of 5 ml/kg.

### Estrous cycle determination

The phase of the estrous cycle was assessed by histological examination of cells extracted by vaginal lavage (Byers, Wiles, Dunn, & Taft, 2012). Briefly, mice were gently restrained and 20 μl of saline were flushed 5 times into the vagina. The resulting fluid was placed onto gelatinized slides, stained with methylene blue and observed at 40X magnification under a light microscope (DM6000 B, Leica Biosystems, Nussloch, Germany).

### Surgical induction of endometriosis

Endometriotic lesions were surgically-induced as previously described (Somigliana et al., 1999), with some modifications. Briefly, uterine horns from donor mice at diestrus were excised, opened longitudinally and biopsied into 4 pieces (2 x 2 mm). Recipient mice were anesthetized with vaporized isoflurane in oxygen (4% V/V for induction; 2.5% V/V for maintenance) and a midline incision of 1 cm was made to expose the pelvic compartment. Endometriosis mice had 4 uterine fragments sutured to the parietal peritoneum, whereas sham-operated mice received 4 similar-sized fragments of abdominal fat. Transplanted tissues and abdominal muscle and skin were stitched using 6-0 black silk (8065195601, Alcon® Cusi S.A., Barcelona, Spain).

### Experimental protocols

The nociceptive, affective and cognitive manifestations associated to the presence of ectopic endometrium were determined in a first experiment. After the measurement of baseline mechanical sensitivity (day −1), endometriosis or sham surgery was performed (day 0), and nociceptive responses were assessed again 7, 14, 21 and 28 days after surgery. Anxiety-like behavior and cognitive performance were evaluated on days 23 and 27, respectively. At the end of the experimental sequence (day 32), mice were euthanized by cervical dislocation for sample collection.

A second experiment was conducted to obtain the ED50 of acute THC administration for the alleviation of mechanical hypersensitivity. Endometriosis and sham mice were tested in the von Frey assay after administration of different doses of THC (1.25, 2, 2.5 and 5 mg/kg) or vehicle. Measurements were done 45 min after subcutaneous administration of THC or vehicle at time points in which endometriotic lesions and pelvic hypersensitivity were fully developed (days 33-41).

The effects of chronic THC or vehicle were evaluated in endometriosis and sham mice in a third experiment. Chronic treatment with THC (2 mg/kg) or vehicle administered once a day (9 AM) started on day 1 after surgery and lasted until day 32. Behavioral measures were conducted as in the first experiment. Mice were tested on the nociceptive paradigm 45 min after drug or vehicle administration and on the anxiety-like and memory tests 6 h after administration. Mice were euthanized on day 32 by cervical dislocation for sample collection.

### Nociceptive behavior

Mechanical sensitivity was quantified by measuring the responses to von Frey filament stimulation of the pelvic area. Von Frey filaments (1.65, 2.36, 2.44, 2.83, 3.22 and 3.61 corresponding to 0.008, 0.02, 0.04, 0.07, 0.16 and 0.4 g; Bioseb, Pinellas Park, FL, USA) were applied in increasing order of force, 10 times each, for 1-2 sec, with an inter-stimulus interval of 5-10 sec. Sharp retraction of abdomen, immediate licking, jumping and scratching of the site of application were considered positive responses. The area under the curve of the data representing the frequency of responses versus the force of the von Frey filaments was calculated using the linear trapezoidal rule for each time point.

### Anxiety-like behavior

The elevated plus maze test was used to evaluate anxiety-like behavior in a black Plexiglas apparatus consisting of 4 arms (29 cm long x 5 cm wide), 2 open and 2 closed, set in cross from a neutral central square (5 x 5 cm) elevated 40 cm above the floor. Light intensity in the open and closed arms was 45 and 5 lux, respectively. Mice were placed in the central square facing one of the open arms and tested for 5 min. The percentage of entries and time spent in the open and closed arms of the maze was determined as a measure of anxiety-like behavior.

### Cognitive behavior

The novel object recognition task was assayed in a V-shaped maze to measure cognitive performance (Puighermanal et al., 2009). On the first day, mice were habituated for 9 min to the maze. On the second day, mice were placed again in the maze for 9 min and 2 identical objects were presented at the ends of the arms of the maze. Twenty-four h later, one of the familiar objects was replaced with a novel one and mice were placed back in the maze for 9 min. The time spent exploring each object (novel and familiar) was recorded and a discrimination index (DI) was calculated as the difference between the time spent exploring the novel and the familiar object, divided by the total time exploring the 2 objects.

### Sample harvesting and tissue preparation

Endometriotic lesions, uterine horns and ovaries were harvested from each mouse and fixed in 4% paraformaldehyde in phosphate buffered saline (PBS) for 4 h and cryoprotected in 30% sucrose with 0.1% sodium azide for 6 days. Samples were embedded in molds filled with optimal cutting temperature compound (4583, Sakura Finetek Europe B.V., Alphen aan den Rijn, The Netherlands) and stored at −80°C until use. Medial prefrontal cortices and hippocampi were fresh-frozen on dry ice immediately after euthanasia and stored at −80°C until use.

### Histology and immunostaining

Endometriotic lesions and uteri were serially sectioned at 20 μm with a cryostat (CM3050, Leica Biosystems, Nussloch, Germany), mounted onto gelatinized slides and stored at −20°C until use. Sections of endometriotic lesions and uteri were stained with hematoxylin and eosin and observed under a Macro Zoom Fluorescence Microscope (MVX10, Olympus, Tokyo, Japan) for assessment of diameter and histological features.

Cyst sections were blocked and permeabilized with 3% normal donkey serum in PBS with 0.3% Triton X-100 for 2 h and incubated overnight with rabbit anti-beta-III tubulin antibody (ab18207, 1:2000, Abcam, Cambridge, United Kingdom) in 3% normal donkey serum in PBS with 0.3% Triton X-100 at 4°C. After washing with PBS, sections were incubated for 1 h at room temperature with anti-rabbit Alexa Fluor A488 antibody (A21206, 1:1000, Thermo Fisher Scientific, Waltham, MA, USA). Slides were washed with PBS and coverslipped with DAPI Fluoromount-G (0100-20, SouthernBiotech, Birmingham, AL, USA) mounting media.

Uterine sections were blocked and permeabilized with 5% normal goat serum in PBS with 0.3% Triton X-100 for 2 h and incubated overnight with rabbit anti-beta-III tubulin antibody (ab18207, 1:2000, Abcam) in 5% normal goat serum in PBS with 0.3% Triton X-100 at 4°C. After washing with PBS, sections were incubated for 1 h at room temperature with anti-rabbit Alexa Fluor A555 antibody (ab150078, 1:1000, Abcam, Cambridge, United Kingdom). Slides were washed with PBS and coverslipped with DAPI Fluoromount-G (0100-20, SouthernBiotech).

### Image analysis

Images of immunostained sections of cysts and uteri were captured with the X2 objective of a Macro Zoom Fluorescence Microscope (MVX10, Olympus, Shinjuku, Tokyo, Japan) and processed and quantified using the NIH Image J software. A blind observer converted from 4 to 8 images per animal into negative black-and-white images and the threshold was manually adjusted. Images were then dilated, skeletonized and the mean percentage of immunoreactive area was obtained by running the “Analyze particles” function.

### Ovarian follicle counting

Sections of ovaries were stained with hematoxylin and eosin and observed under an upright microscope (DM6000 B, Leica Biosystems). The number of pre-antral and antral follicles was determined in every 9 sections. Only follicles containing an oocyte were counted and the total number of follicles was estimated by multiplying the raw counts by 9 according to published criteria (Myers, Britt, Wreford, Ebling, & Kerr, 2004). The number of corpora lutea was determined by direct counting of every 18 sections according to the average corpus luteum diameter (Numazawa & Kawashima, 1982).

### Gene expression analysis by RT-PCR

Total RNA was isolated from frozen medial prefrontal cortices and hippocampi with a RNAqueous™-Micro Total RNA Isolation Kit (AM1931, Invitrogen, Waltham, MA, USA) and subsequently reverse-transcribed to cDNA with a High Capacity cDNA Reverse Transcription Kit (4368814, Applied Biosystems, Foster City, CA, USA) according to the manufacturer’s instructions. RT-PCR was carried out in triplicate with a QuantStudio 12K Flex Real-Time PCR System (4471134, Applied Biosystems, Foster City, CA, USA) using the SYBR Green PCR Master Mix (04707516001, Roche, Basel, Switzerland). Relative expression values were normalized to the expression value of *GAPDH* (control gene). The following specific primers were used: 5’-CGTGGAGTCCGCTTTACAG-3’ (*Ptgs2* forward); 5’-CCTTCGTGAGCAGAGTCCTG-3’ (*Ptgs2* reverse), 5’-GAAGTTGACGGACCCCAAAA-3’ (*Il1b* forward); 5’-TGATGTGCTGCTGCGAGATT-3’ (*Il1b* reverse); 5’-AGAGGGATGCTGCCCTTACC-3’ (*GAPDH* forward); 5’-ATCCGTTCACACCGACCTTC-3’ (*GADPH* reverse). Data for each target gene were analyzed by the 2^−ΔΔCt^ method after normalization to the endogenous control *GAPDH* (Livak & Schmittgen, 2001).

### Statistical analysis

Data obtained with the nociception model were analyzed using one-way repeated measures ANOVA (surgery as between-subject factor) or two-way repeated measures ANOVA (surgery and treatment as between-subject factors). Dose-response curve was fitted and ED50 determined using GraphPad Prism 7 (San Diego, CA, USA). Data obtained with the elevated plus maze test, novel object recognition task, histology, immunostaining and ovarian follicle counting were analyzed using a Student t-test (surgery) or a two-way ANOVA (surgery and treatment). Post hoc Bonferroni analysis was performed after ANOVA when appropriate. Since the equality of variances could not be confirmed for all gene expression data, the nonparametric Kruskal-Wallis test was used in this case, followed by Mann Whitney U when appropriate. Data are expressed as individual data points and mean ± SEM, and statistical analyses were performed using IBM SPSS 18 software (Chicago, IL, USA). The differences were considered statistically significant when the p value was below 0.05.

## Author contributions

All authors listed above have contributed sufficiently to be included as authors. A.E.L. conducted the behavioral and molecular experiments, performed immunohistochemistry and microscopy, and wrote the manuscript. J.A. performed immunohistochemistry and microscopy. D.C. conceptualized and supervised the project, participated in the experimental design and wrote the manuscript. R.M. conceptualized, supervised and funded the project, participated in the experimental design and wrote the manuscript. All the authors have revised the work critically for important intellectual content and approved the final version to be published.

## Acknowledgements

Authors thank Mercè Vilaró Blay and Berta Güell Villena for their technical help.

## Data availability

Authors declare that all data supporting the findings of this study are available within the manuscript and its source data files.

---

**Figure 1- figure supplement 1.**
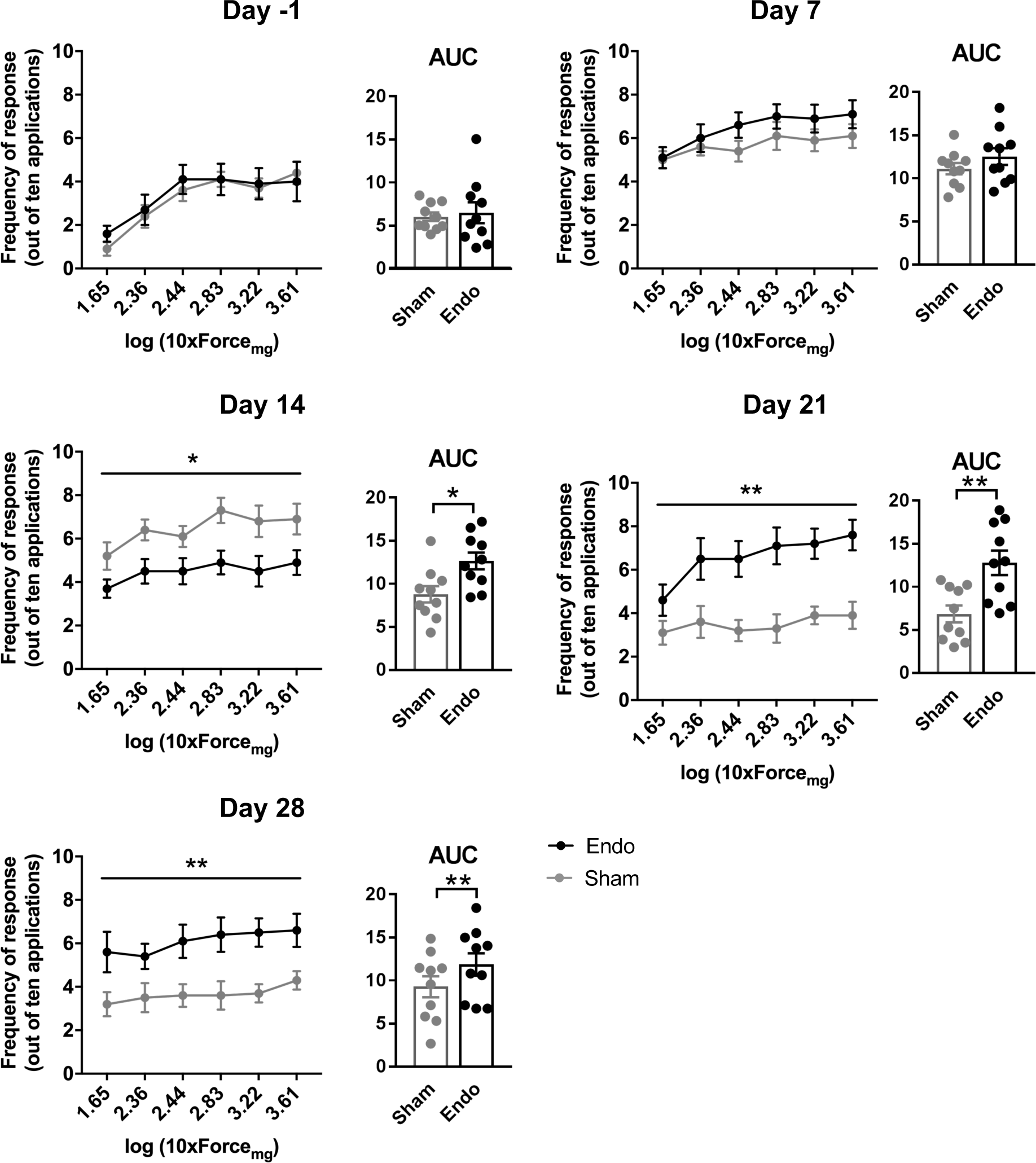
Nociceptive responses to mechanical stimulation with von Frey filaments. Significantly higher frequency of responses and AUC was observed in endometriosis mice when compared to sham mice on days 14, 21 and 28 after the surgery. For each day, left panel is frequency of response to each von Frey filament and right panel is the corresponding AUC. Error bars are mean ± SEM. For each day, one-way repeated measures ANOVA (left panels) and Student t-test (right panels). *p<0.05, **p<0.01 vs sham. Endo, endometriosis; AUC, area under the curve. Figure 1 - Source Data.

**Figure 1 – figure supplement 2.**
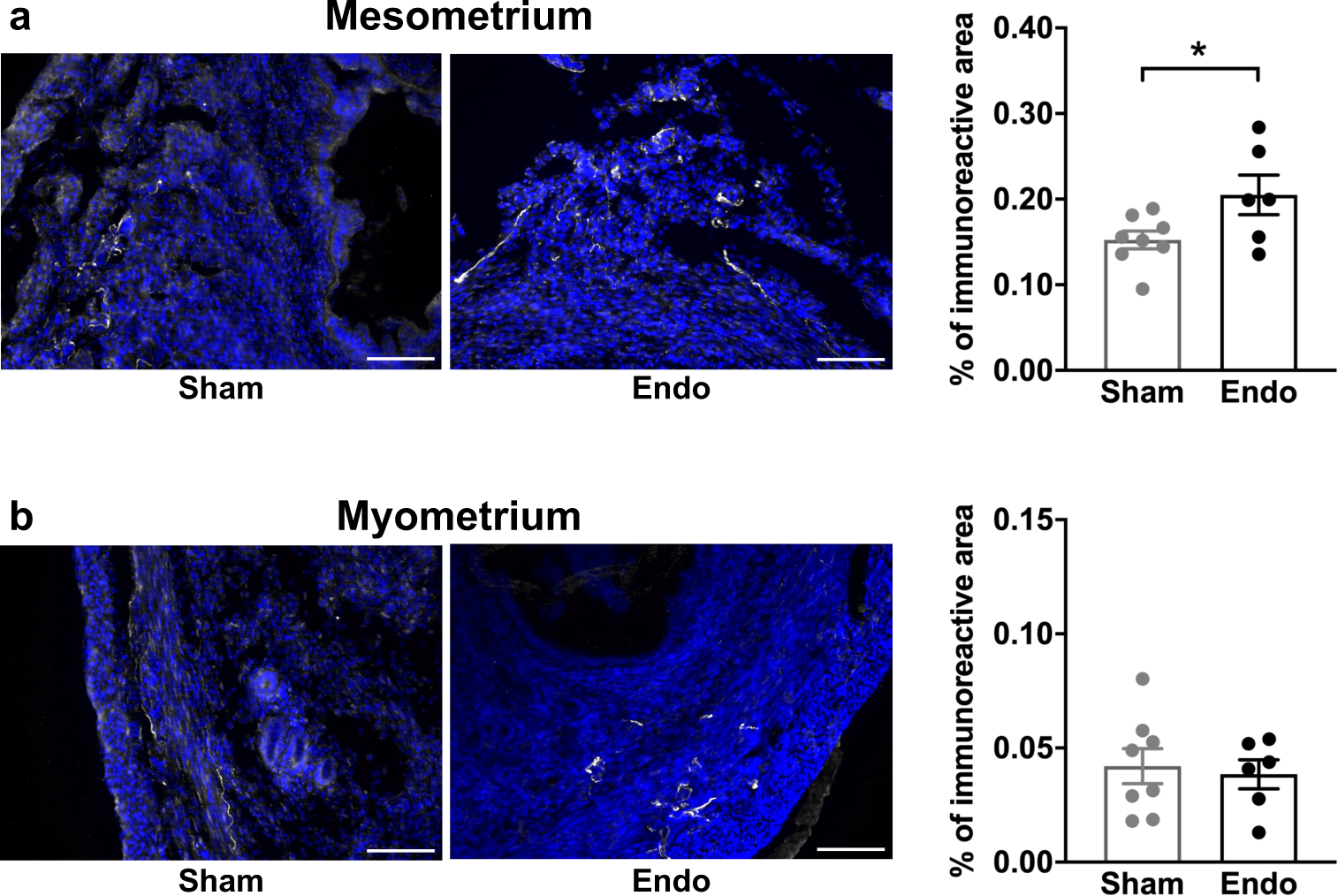
Density of beta-III tubulin-labeled fibers in uteri of endometriosis and sham mice. (**a**) The percentage of immunoreactive area of the mesometrial aspect of the uterus was higher in endometriosis mice. (**b**) The percentage of immunoreactive area of myometrium did not differ between groups. Blue is DAPI and white is β-III tubulin. Scale bar = 100 μm. Error bars are mean ± SEM. Student t-test. *p<0.05 vs sham. Endo, endometriosis. Figure 1 - Source Data.

**Figure 3 – figure supplement 1.**
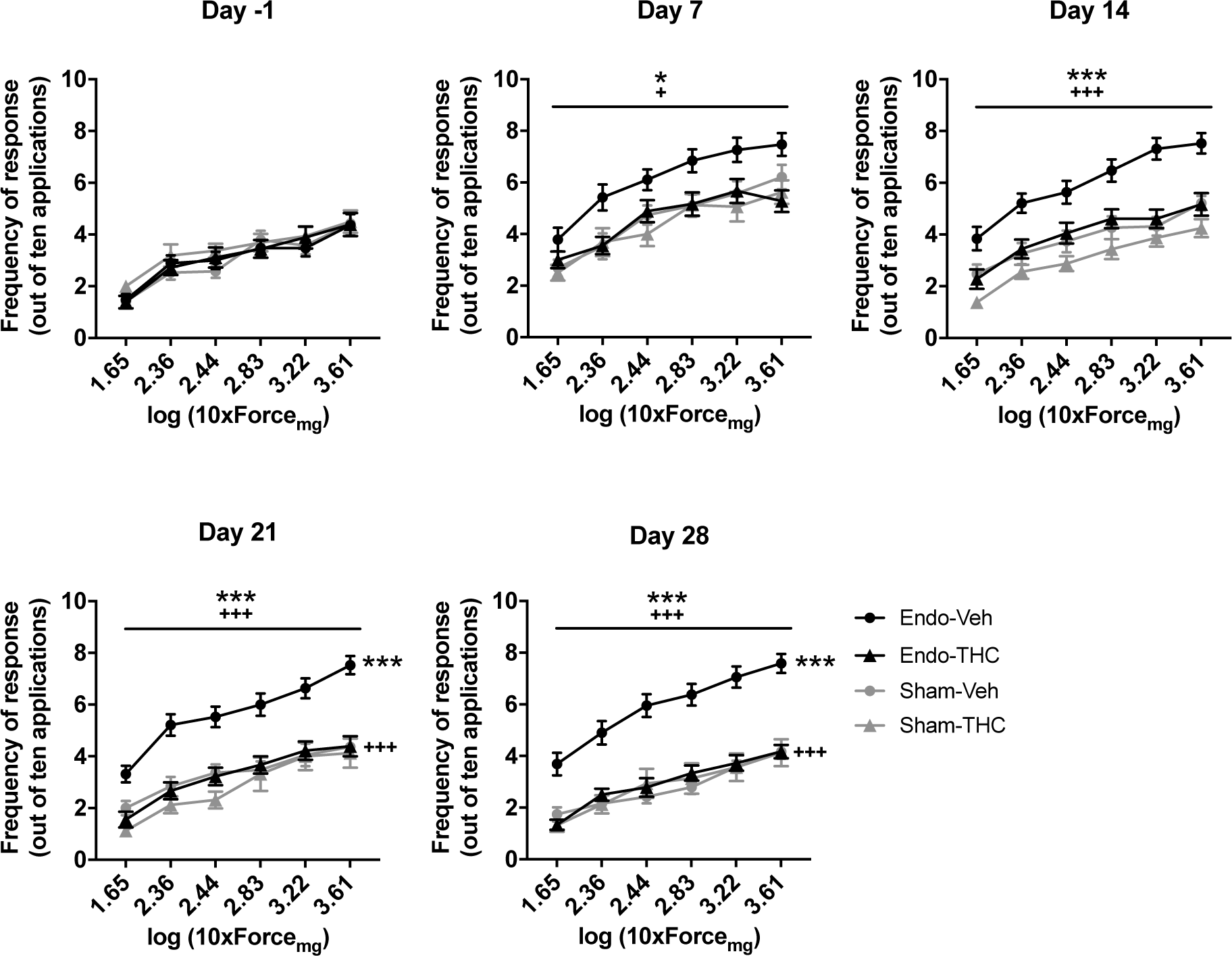
Effect of chronic THC treatment on nociceptive responses to mechanical stimulation. Endometriosis mice treated with vehicle showed higher frequency of response than endometriosis mice treated with THC and sham mice treated with vehicle. Error bars are mean ± SEM. Two-way repeated measures ANOVA + Bonferroni. *p<0.05, ***p<0.001 vs sham. +p<0.05, +++p<0.001 vs vehicle. Endo, endometriosis; Veh, vehicle; THC, Δ9-tetrahydrocannabinol. Figure 3 - Source Data.

**Figure 3 – figure supplement 2.**
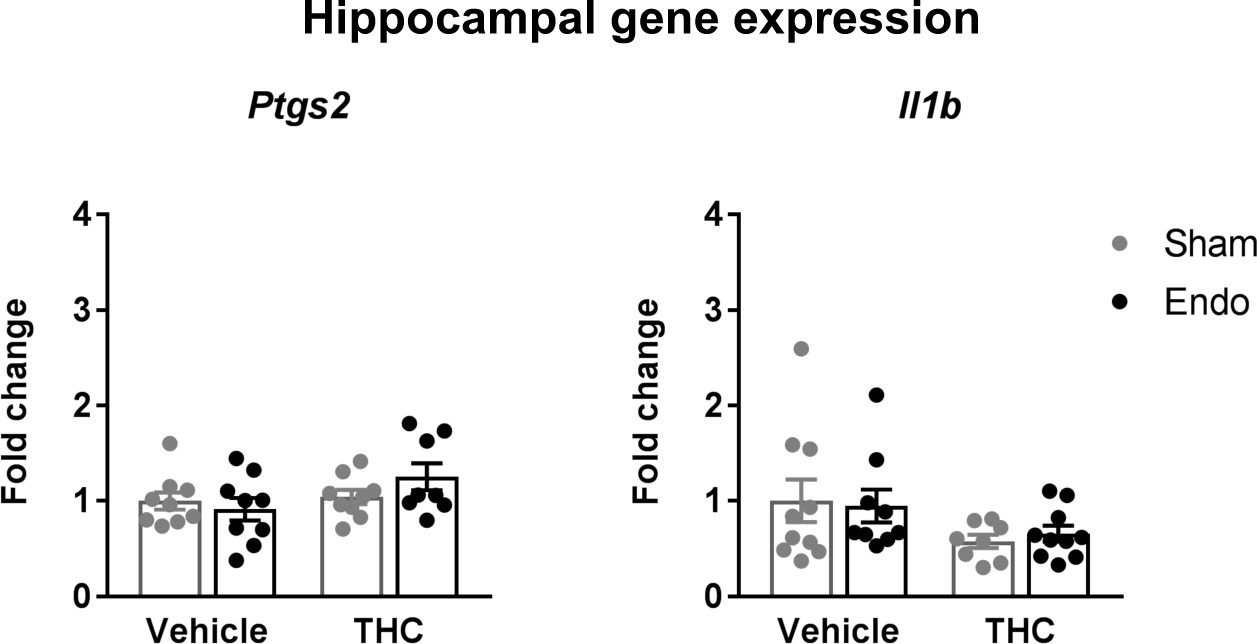
Gene expression levels of inflammatory markers in the hippocampus. mRNA levels of the genes coding for interleukin 1b (*Il1b*) and cyclooxygenase 2 (*Ptgs2*) remained unaltered. Error bars are mean ± SEM. Kruskal-Wallis. Endo, endometriosis; THC, Δ9-tetrahydrocannabinol. Figure 3 - Source Data.

**Figure 4 – figure supplement 1.**
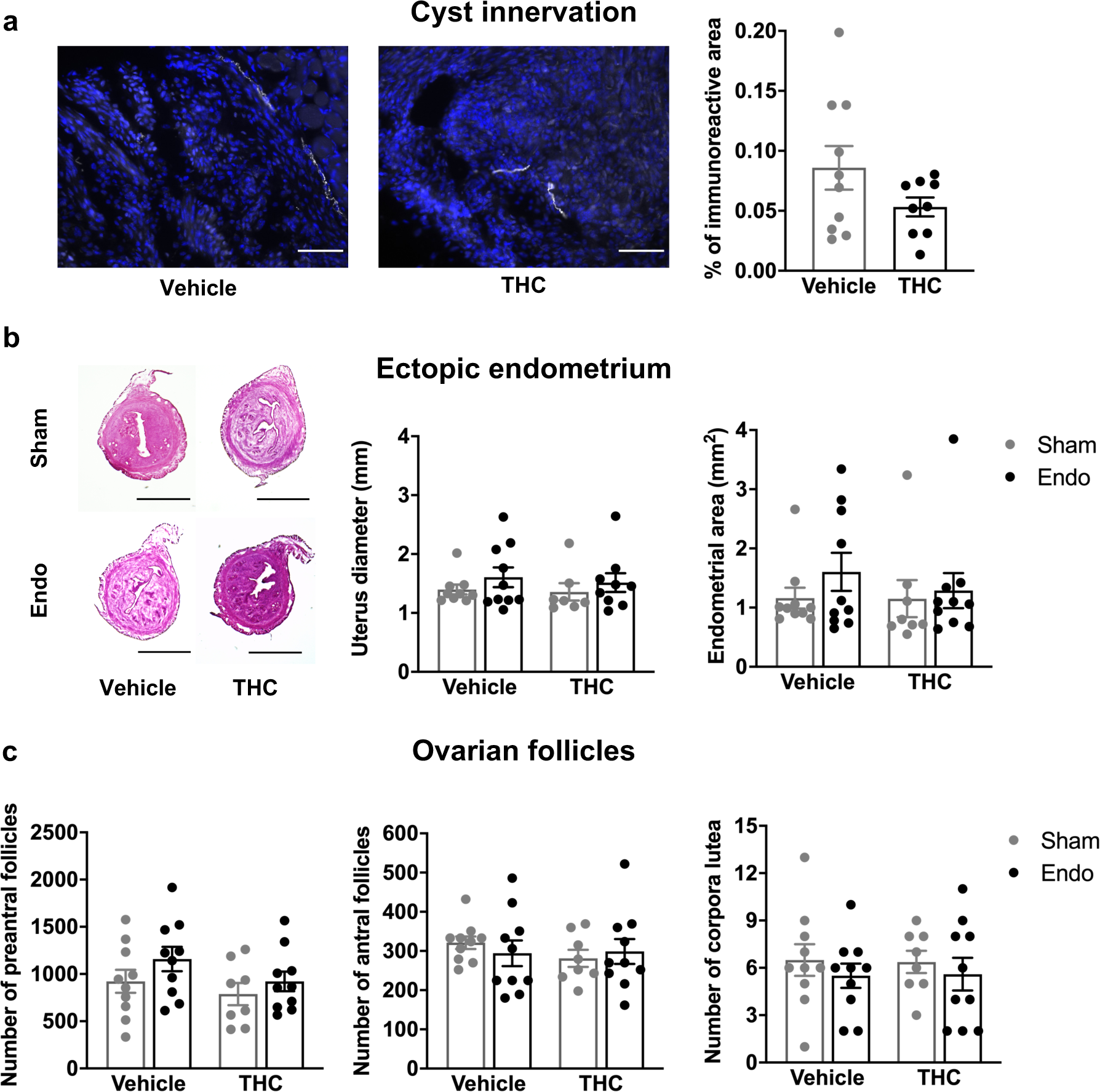
Histological features of reproductive tissues after chronic THC treatment. (**a**) Cyst innervation was unaffected by THC. Blue is DAPI and white is β-III tubulin. Scale bar = 100 μm. (**b**) Uterine diameter and area of endometrial tissue were similar among the groups. Scale bar = 1 mm. (**c**) Number of preantral follicles, antral follicles and corpora lutea were unchanged after THC. Error bars are mean ± SEM. Student t-test (a) and two-way ANOVA (b and c). Endo, endometriosis; THC, Δ9-tetrahydrocannabinol. Figure 4 - Source Data.

## List of source data files

- Figure 1 - Source Data. Effects of ectopic endometrium.
- Figure 2 - Source Data. Acute THC effects.
- Figure 3 - Source Data. Effects of repeated THC on behavioral alterations.
- Figure 4 - Source Data. Effects of repeated THC on histopathological features.

